# Bone marrow stromal cells induce an ALDH^+^ stem cell-like phenotype in AML cells through TGF-β-p38-ALDH2 pathway

**DOI:** 10.1101/2020.07.10.197178

**Authors:** Bin Yuan, Fouad El Dana, Stanley Ly, Yuanqing Yan, Vivian Ruvolo, Elizabeth J. Shpall, Marina Konopleva, Michael Andreeff, V. Lokesh Battula

**Affiliations:** Section of Molecular Hematology and Therapy, Department of Leukemia, Houston, Texas, USA; Department of Biostatistics, Houston, Texas, USA; Department of Stem Cell Transplantation and Cellular Therapy, The University of Texas MD Anderson Cancer Center, Houston, Texas, USA

**Keywords:** Mesenchymal stromal cells, AML, ALDH, ALDH2, TGF-β, non-canonical pathway, p38, bone marrow microenvironment

## Abstract

Mesenchymal stromal cells (MSCs) in the bone marrow (BM) microenvironment have been shown to induce chemotherapy resistance in acute myeloid leukemia (AML) cells, but the mechanism is not clear. We hypothesized that stromal cells induce a stem-like phenotype in AML cells, thereby promoting tumorigenicity and chemotherapy resistance. We found that aldehyde dehydrogenase (ALDH), an enzyme that is highly expressed in hematopoietic as well as leukemic stem cells was dramatically activated in AML cells co-cultured with BM-MSCs mainly through upregulation of a specific isoform, ALDH2. Mechanistic studies revealed that stroma-derived TGF-β1 induced an ALDH^+^ phenotype in AML cells via the non-canonical TGF-β pathway through p38 activation. Inhibition of ALDH2 using specific inhibitors significantly inhibited BM-MSC-induced ALDH activity and sensitized AML cells to chemotherapy. Collectively, our data indicate that BM stroma induces a stem-like phenotype in AML cells through the non-canonical TGF-β pathway. Inhibition of ALDH2 sensitizes AML cells to chemotherapy.

**Impact Statement:** Currently there is no standard therapy for AML. In this study we identified the mechanism of chemotherapy resistance in AML cells and discovered TGF-β-p38-ALDH2 signaling pathway as a therapeutic target.

## Introduction

The bone marrow microenvironment (BME) contributes to Acute Myeloid Leukemia (AML) growth and chemotherapy resistance (1, 2). Mesenchymal stromal cells (MSCs) in the bone marrow (BM) are critical for growth induction and anti-apoptotic signaling in AML (3). However, the mechanisms of stroma-mediated AML growth and chemotherapy resistance are not clear. As leukemogenesis and chemotherapy resistance are characteristics of AML stem cells, we hypothesized that the BME induces a stem cell-like phenotype in AML cells.

Several signaling pathways contribute to chemotherapy resistance in AML through induction of a high-mesenchymal stem-like cell state (4). Among them, transforming growth factor-β (TGF-β)-mediated canonical and non-canonical pathways have been well-characterized in AML cells (5–7). Inhibition of TGF-β signaling using small molecule inhibitors or receptor-blocking antibodies inhibited leukemia growth and sensitized AML cells to chemotherapy (5). TGF-β signaling has cell type–specific effects and has been involved in the induction of a stem cell–like phenotype in solid tumors (8–11).

Aldehyde dehydrogenase (ALDH) is an enzyme involved in oxidizing toxic aldehydes into neutral acids (12). ALDH activity is increased in hematopoietic stem cells and leukemia stem cells (13, 14). ALDH-positive (ALDH^+^) leukemia cells have higher tumorigenicity and chemotherapy resistance compared to ALDH-negative cells (14, 15). Additionally, high ALDH activity at diagnosis predicts relapse in a subset of AML patients (16). Among the 19 ALDH isoforms identified in humans, the most prominent are ALDH1 isoforms (ALDH1A1-3, ALDHB1, and ALDH1L1&2) and the ALDH2 isoform. ALDH1 family isoforms are located in the cytoplasm, whereas ALDH2 is located in the mitochondria (17, 18).

In this report, we investigated the effect of BM stromal cells on AML cells, signaling pathways activated, and therapeutic targets that contribute to chemotherapy resistance in AML cells. We identified specific inhibitors that could be used in combination with standard chemotherapy for treatment of AML patients.

## Methods

### Cell culture of primary MSCs, leukemia cell lines, and patient samples

HL-60 cells were purchased from ATCC^®^ and MOLM-13 cells were obtained from the MD Anderson Cell Line core facility. Both cell lines were cultured in RPMI (Media Tech, Inc., Manassas, VA) with 10% fetal bovine serum (FBS) and 1% penicillin/streptomycin. OCI-AML3 cells were a kind gift from Dr. Mark Minden at Ontario Cancer Institute, Toronto, Canada. Human MSCs were isolated from BM harvested from healthy patient donors for use in allogeneic stem cell transplantation. All donors were consented. This study was performed according to a protocol approved by the institutional review board of the University of Texas MD Anderson Cancer Center. Tests for *Mycoplasma* contamination of MSCs are performed in our laboratory every 4-6 months. Human MSCs were cultured in MSC Growth Medium 2 (Cat# C-28009, PromoCell^®^, Heidelberg, Germany) with 1% penicillin/streptomycin. Peripheral blood mononuclear cells of patients were cultured in RPMI containing 10% FBS and 1% penicillin/streptomycin.

### Flow cytometry assay

Flow cytometry analysis of AML cells cultured alone or co-cultured with MSCs was performed as described before (19). The cells were incubated with fluorochrome-conjugated antibodies for 20 minutes. The antibody conjugates used were anti-CD45 conjugated with APC (Cat# 304038, BioLegend^®^, San Diego, CA) and anti-CD90 conjugated with APC/Alexafluor 750 (Cat# B36121, Becton Dickinson Biosciences, Franklin Lakes, NJ). Cells were then washed with phosphate buffered saline (PBS) containing 0.5 μg/mL 4′, 6-diamino-2-phenylindole (DAPI; Cat# D1306, ThermoFisher Scientific, Waltham, MA) to exclude dead cells and analyzed on an LSR-II Flow Cytometer (BD Biosciences). Twenty thousand events were acquired for each sample. Data were analyzed by FlowJo software (FlowJo LLC, Ashland, Oregon).

### ALDH activity assay

ALDH assay was performed as per the ALDEFLUOR™ kit (Cat #01700, STEMCELL^TM^ Technologies, Vancouver, Canada) manufacturer’s instructions. The cells were washed once with ALDEFLUOR™ buffer and stained with CD90 (to exclude MSCs during analysis) at 4°C for 30 minutes, then re-suspended with 0.3 mL ALDEFLUOR™ buffer and analyzed by flow cytometry. To determine the effect of ALDH2 inhibition on total ALDH activity, the cells were treated with the ALDH2 inhibitors diadzin (Cat# CS-4237, ChemScene, Monmouth Junction, NJ) and CVT-10216 (Cat # SML1366-5mg, Sigma Aldrich). ALDH activity was measured as described above.

### Protein analysis by western blotting

Cells were lysed in RIPA buffer at 3×10^5^/50 μL density. Protein concentrations were determined using Bradford protein assay. Laemmli buffer was added to protein lysates at a 1:1 ratio. The lysates were loaded onto 4–15% Mini-PROTEAN^®^ TGX™ Precast Protein Gels (Cat# 4561086, Bio-Rad, Hercules, CA), and proteins were subsequently transferred onto a polyvinylidene fluoride (PVDF) membrane. The membrane was blocked with 5% milk in PBS-T (0.05% Tween-20 in PBS) to prevent nonspecific binding of antibodies. Primary antibody incubation was performed in PBS-T with 1% milk at 4°C overnight (refer to Supplementary Table 1 for list of primary antibodies used). IRDye^®^ 680RD donkey anti-rabbit IgG or IRDye^®^ 800CW goat anti-mouse IgG (LI-COR Biosciences^®^, Lincoln, NE) was incubated with the membranes for 1 hour at room temperature in PBS-T with 1% milk. The membranes were washed 3 times with PBS-T and scanned using an Odyssey western blot scanner (LI-COR Biosciences^®^). All protein quantification was performed using LI-COR image analysis software.

### shRNA knockdown of TGF-β1 expression

Lentiviral-mediated short-hairpin RNA (shRNA) was used for stable knockdown of TGF-β1 in human BM-derived MSCs. shRNA lentiviral vectors (NM_000660, XM_011527242; Clone ID: TRCN0000003318; Sequencing Primer: 5’ - AAACCCAGGGCTGCCTTGGAAAAG - 3’; Vector Map: pLKO.1) were purchased from GE Healthcare Dharmacon, Inc. (Lafayette, CO). Lentiviral pLKO.1 Empty Vector (Cat# RHS4080, GE-Dharmacon) was used as control. HEK293T cells were transfected with each TGF-β1 shRNA construct along with packaging vectors pMD2.G (0.5 μg) and psPAX2 (1.5 μg) (Addgene, Inc., Watertown, MA) using Jet Prime Reagent (Polyplus-transfection^®^, Illkirch-Graffenstaden, France) according to manufacturer’s guidelines. The medium containing lentivirus was collected 72 hours after transfection and incubated with BM-MSC cells for 24 hours. The transduced cells were selected using puromycin (0.5 μg/mL) for 3 days, and TGF-β1 mRNA and protein knockdown efficacy was determined by quantitative polymerase chain reaction (qPCR) or western blotting, respectively. The plasmid TRCN0000003318 (RHS4533-EG7040, GE-Dharmacon) provided the best knockdown efficacy.

### Total RNA isolation and gene expression by real-time PCR

Five million cells were lysed in Trizol reagent overnight at −80°C, total RNA was isolated by ethanol precipitation, and real-time PCR (RT-PCR) was performed with a QuantStudio3 (Applied Biosystems^®^, Foster City, CA) instrument using TaqMan Fast Universal PCR Master Mix (Applied Biosystems^®^) as described before (20). All samples were run in triplicates. The relative fold increase of specific RNA was calculated by the comparative cycle of threshold detection method, and values were normalized to Glyceraldehyde 3-phosphate dehydrogenase (GAPDH). Fold changes in gene expression were calculated using the 2-ddC method. All primer pairs for human samples were purchased from ThermoFisher Scientific (Supplementary Table 2).

### AML cell separation by fluorescence-activated cell sorting

OCI-AML3 and BM-MSC cells were seeded at a 5:1 ratio in RPMI medium with 10% FBS and cultured for 3-5 days. On the day of cell separation, OCI-AML3 cells were collected. After a single wash with PBS, cells were stained with APC-Cy7-conjugated anti-CD90 (Cat # 16699531, BD Biosciences) and FITC-conjugated anti-CD45 (Cat # 304038, BioLegend^®^) antibodies. Cells were washed with PBS containing DAPI solution, then subjected to fluorescence-activated cell sorting (FACS) using a BD FACS Aria-II cell sorter (BD Biosciences). To isolate AML cells, the cells were gated on a CD45^+^CD90^−^ fraction using FACSDiva software.

### Gene expression analysis by RNA sequencing

Samples were sequenced on the HiSeq Sequencing System by the Sequencing and Microarray Core facility at MD Anderson Cancer Center. Sequence reads were mapped to human genomics (build hg19) with bowtie2 aligner using RSEM software. The EdgeR package in R software was used to compare the differential expression between OCI-AML3 cells cultured alone or with BM-MSCs. Genes with adjusted p values less than 0.05 and absolute fold changes larger than 2 were considered significant. To investigate the relationship between the significant genes differentially expressed in the presence of BM-MSCs, we used AML expression data obtained from the TCGA dataset. A scatterplot with log2 fold change from expression in co-culture and TCGA correlation coefficients was plotted using the cBioPortal data analysis tool. The array data has been deposited in the Gene Expression Omnibus (GEO) identified by the accession number GSE152996.

### p38 MAPK inhibition

The p38 MAPK inhibitor SB-203580 was purchased from R#D Systems (Cat# 1202, Minneapolis, MN). OCI-AML3 cells were cultured with or without BM-MSCs and treated with p38 MAPK inhibitor (SB-203580; 2 μM) for 3 days. Similarly, OCI-AML3 cells were treated with p38 MAPK inhibitor in the presence or absence of recombinant TGF-β1 (5 ng/mL) and ALDH activity was measured. To determine the effect of p38 inhibition on ALDH2 expression, OCI-AML3 cells cultured with or without BM-MSCs were treated with SB-203580 at 2 μM for 3 days, and ALDH2 expression was measured by western blotting.

### ALDH2 inhibition in combination with standard chemotherapy

To assess the combined effect of ALDH2 inhibition and chemotherapy on AML cells, OCI-AML3 cells were cultured at 0.25 million cells/mL density with or without BM-MSCs (100 000 cells). AML cells were treated with cytarabine (Ara-C; 2.5 μM; obtained from the MD Anderson pharmacy), alone or in combination with CVT-10216 (5 μM), for 48 hours. Cells were stained with Annexin V and analyzed on an LSR-II flow cytometer.

### Leukemia Engraftment and growth rate

To investigate the effect of stromal cells on leukemia engraftment and growth, we implanted one million Molm13 cells expressing firefly luciferase and GFP subcutaneously, with or without 1 million BM-MSCs and 100 μL Matrigel, in Nonobese Diabetic/ Severe Combined Immunodeficiency (NOD/SCID) mice. Leukemia engraftment and growth rate assessment was performed at 1 and 2 weeks as previously described (19).

### Animal study approval

All animal experiments were in compliance with a protocol approved by the MD Anderson Cancer Center Institutional Animal Care and Use Committee.

### Statistical analyses

For survival analysis, we used Kaplan-Meier estimator to estimate the survival function and log-rank test to evaluate the statistical significance. To compare the difference two independent groups, we used Mann-Whitney U-test or Student’s T-test to examine the statistical significance. For the comparison with two groups with paired data, paired Student’s T-test was used. We performed one way ANOVA with Tukey’s HSD post hoc test to test the significance in the comparison with more than 2 groups. A linear model with interaction term was also used to evaluate the significance with more than two factors in the experiment. A P-value less than 0.05 was regarded as statistically significant.

## Results

### BM-MSCs induce leukemia growth in vivo by inducing ALDH activity in AML cells

BM stroma contributes to AML progression and chemotherapy resistance (3, 19, 21). To investigate the effect of stromal cells on AML growth, we implanted Molm13 cells, with or without BM-MSCs, subcutaneously into NOD/SCID mice. Bioluminescence imaging revealed that Molm13 cells implanted together with BM-MSCs grew 8-fold faster than Molm13 cells implanted alone, suggesting that BM-MSCs support AML cell growth (Fig. 1A, 1B).

**Figure 1.**
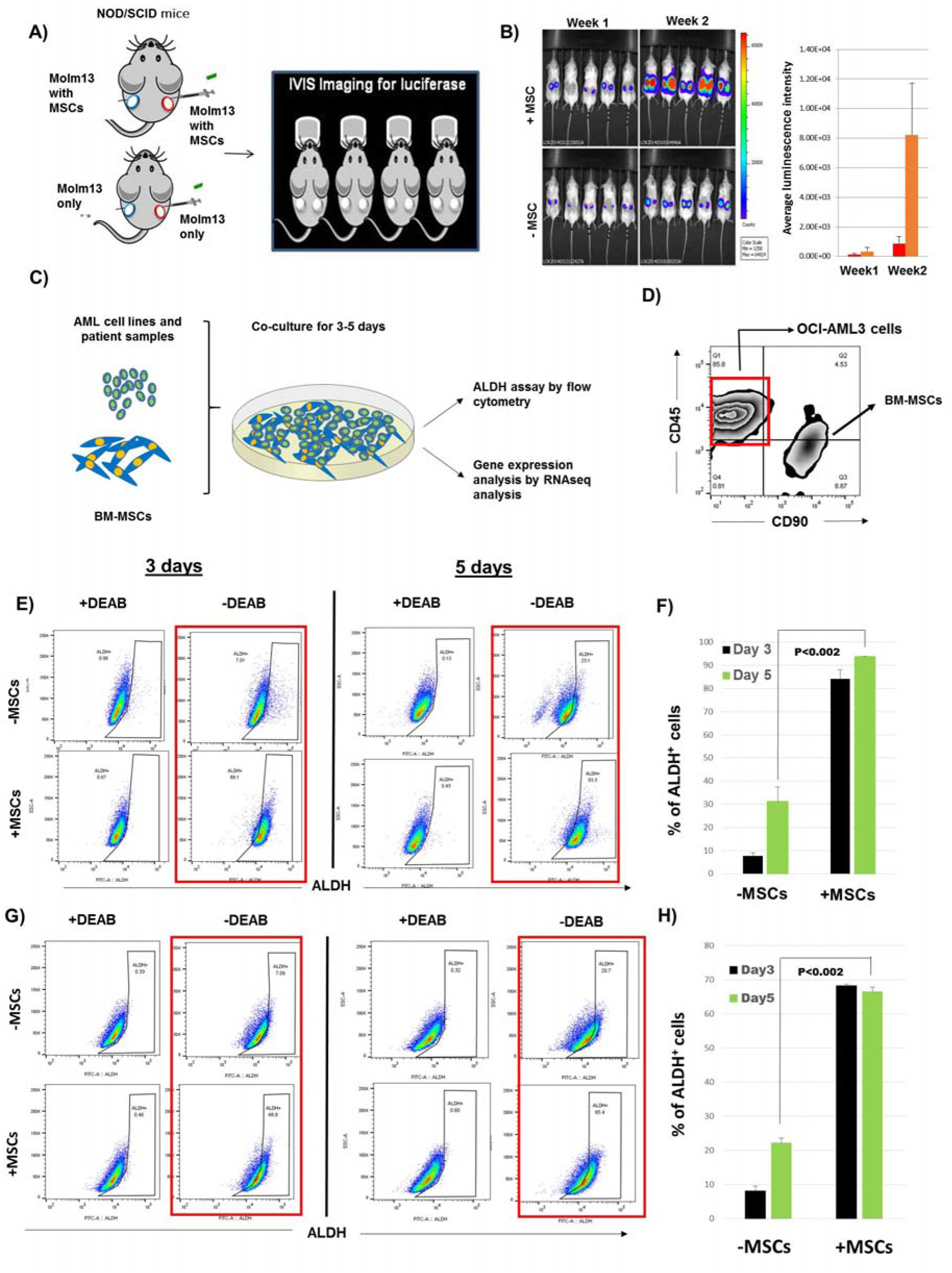
BM-MSCs induce ALDH activity in AML cells and enhance engraftment in mice. **(A, B)** One million firefly luciferase-expressing AML cells (Molm13) were implanted subcutaneously, with or without 1 million BM-MSCs and 100 μL of Matrigel, in NOD/SCID mice. Bioluminescence imaging was performed at 1 and 2 weeks to check leukemia engraftment and growth rate. **(C)** AML cell lines and patient samples were cultured with or without BM-MSCs for 3-5 days. ALDH activity in AML cells was measured using ALDEFLUOR^®^ assay. Gene expression analysis was performed by RNA sequencing. **(D)** Fluorescence-activated cell sorting of OCI-AML3 cells was performed based on phenotype (CD45^+^, CD90^-^) to distinguish them from BM-MSCs (CD90^+^, CD45^-^). **(E)** OCI-AML3 cells were cultured with or without BM-MSCs for 3 or 5 days. Cells were stained with ALDEFLUOR^®^, CD45, and CD90. During FACS analysis, MSCs (CD90^+^, CD45^-^) were gated and ALDH activity was measured in OCI-AML3 cells by flow cytometry. Data were analyzed on FlowJo software. **(F)** Histogram representation of the percentage of ALDH^+^ OCI-AML3 corresponding to the experiment done in **E**. **(G)** The same experiment was done as in **E**, using HL60 AML cells. **(H)** Histogram representation showing the percentage of ALDH^+^ HL60 cells corresponding to the experiment done in **G**. Data are plotted as the mean value with error bars representing standard error. For **B**, **F**, and **H** a linear model with interaction term was used to evaluate the significance. N=3 for each group.

To investigate the effect of BM stromal cells on AML cells, we co-cultured AML cells OCI-AML3 and HL60 with or without BM-MSCs for 3 or 5 days and measured ALDH activity. The percentage of ALDH^+^ cells increased from 31% ± 6% to 94% ± 0.5% when OCI-AML3 cells were co-cultured with BM-MSCs compared to being cultured alone. In HL60 cells, co-culturing with BM-MSCs increased the percentage of ALDH^+^ cells from 22% ± 1% to 76% ± 1% (Fig. 1C-H).

To validate stroma-induced ALDH activity in primary AML cells, we analyzed ALDH activity in peripheral blood and BM samples derived from AML patients and found that the percentage of AML cells varied between patients and did not correlate with age, sex, white blood cell count, or blast percentage (Supplementary Table 3). Next, patient-derived primary AML cells were cultured with or without MSCs for 3 days and ALDH activity was measured in AML cells. We found that co-culture with MSCs significantly induced ALDH activity in AML cells in all 8 patient samples (Fig. 2A, 2B). This indicates that BM-MSCs support AML cell growth in vivo and induce the ALDH^+^ stem cell phenotype in AML cells.

**Figure 2.**
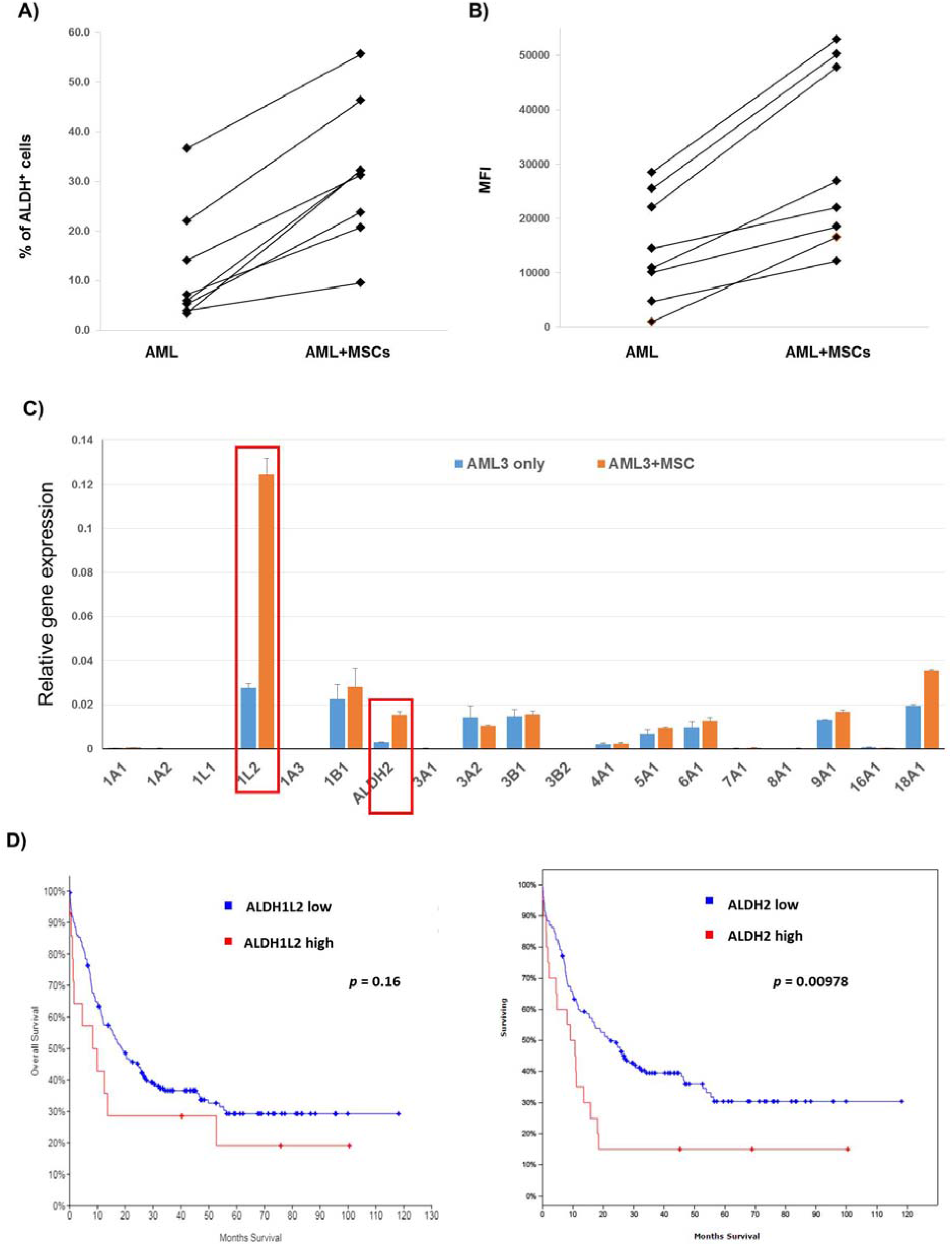
Differential expression of ALDH isoforms in AML cells co-cultured with BM-MSCs. **(A, B)** Patient-derived primary AML cells from bone marrow and peripheral blood samples were cultured with or without MSCs for 3 days and ALDH activity was measured using ALDEFLUOR^®^ assay by flow cytometry. Data show a graphical representation of ALDH mean fluorescence intensity (MFI), and percentage of ALDH^+^ AML patient samples when co-cultured with stromal cells. **(C)** Total RNA was extracted from 5 million AML cells. RT-PCR was performed to analyze gene expression of different ALDH isoforms using primers listed in Supplementary Table 2. All samples were run in triplicate. The relative fold increase of specific RNA was calculated by the comparative cycle of threshold detection method, and values were normalized to GAPDH. Fold changes in gene expression were calculated using the 2-ddC method after normalization to GAPDH. Data are plotted as mean values with error bars representing standard error. **(D)** AML expression data for ALDH isoforms and survival analysis was obtained from the TCGA dataset. A scatterplot with log2 fold change from expression in co-culture and TCGA correlation coefficients was plotted using cBioPortal for cancer genomics data analysis software. For **A** and **B**, paired T-test was used. For **C**, Mann-Whitney U-test/Student’s T-test was used (N=3). For **D**, log-rank test was used to test the significance.

### ALDH isoforms are differentially expressed in AML cells in the presence of stromal cells

To identify the ALDH isoforms responsible for increasing ALDH activity in AML cells in the presence of stromal cells, we performed gene expression analysis of the 19 ALDH isoforms by real-time RT-PCR (primers listed in Supplementary Table 2). We found differential expression of ALDH isoforms in AML cells cultured with or without BM-MSCs. Specifically, ALDH1L2 and ALDH2 expression was upregulated 3- to 5-fold in OCI-AML3 cells co-cultured with BM-MSCs compared to OCI-AML3 cells cultured alone (Fig. 2C). To determine the prognostic significance of these isoforms, we analyzed ALDH1L2 and ALDH2 expression in the TCGA AML dataset, which revealed that ALDH1L2 and ALDH2 are upregulated in 8% and 12% of AML cases, respectively. However, increased expression of ALDH2, but not ALDH1L2, confers a worse prognosis and lower survival rate, suggesting that ALDH2 is a key factor promoting AML disease progression (Fig. 2D).

### TGF-β1-associated gene signature is activated in OCI-AML3 cells co-cultured with BM-MSCs

To investigate the mechanism of stroma-induced ALDH activity in AML cells, we co-cultured OCI-AML3 cells with or without BM-MSCs for 3 days. OCI-AML3 cells were FACS sorted and gene expression analysis was performed by RNA sequencing. Analysis of differentially expressed genes by the Ingenuity^®^ pathway analysis tool revealed activation of a TGF-β1-associated gene signature in OCI-AML3 cells co-cultured with BM-MSCs compared to OCI-AML3 cells cultured alone (Fig. 3A). To validate this, we performed RT-PCR for genes that are differentially regulated by TGF-β1. Genes that are positively regulated by TGF-β1 were upregulated and genes that are negatively regulated by TGF-β1, were downregulated in OCI-AML3 cells co-cultured with MSCs compared to cells cultured alone (Fig. 3 B, 3C). Hence, TGF-β1-regulated transcriptional activity is upregulated in AML cells co-cultured with BM-MSCs. To determine whether TGF-β1 is derived from stromal cells and regulates ALDH activity in AML cells, we knocked down TGF-β1 in BM-MSCs. We found that 2 of 5 shRNA sequences showed highest TGF-β1 knockdown efficacy in BM-MSCs (Supplementary Fig. 2). We then co-cultured OCI-AML3 cells with control (scrambled shRNA) or TGF-β1-knockdown BM-MSCs for 3 days and measured ALDH activity in OCI-AML3 cells. Remarkably, the percentage of ALDH^+^ cells decreased by ~4-fold in OCI-AML3 cells co-cultured with TGF-β1-knockdown BM-MSCs, compared to OCI-AML3 cells cultured with control BM-MSCs (Fig. 3D). Therefore, knockdown of TGF-β1 inhibits stroma-induced ALDH activity, further solidifying the hypothesis that TGF-β1 secreted by stromal cells is directly involved in the increase in ALDH activity in AML cells.

**Figure 3.**
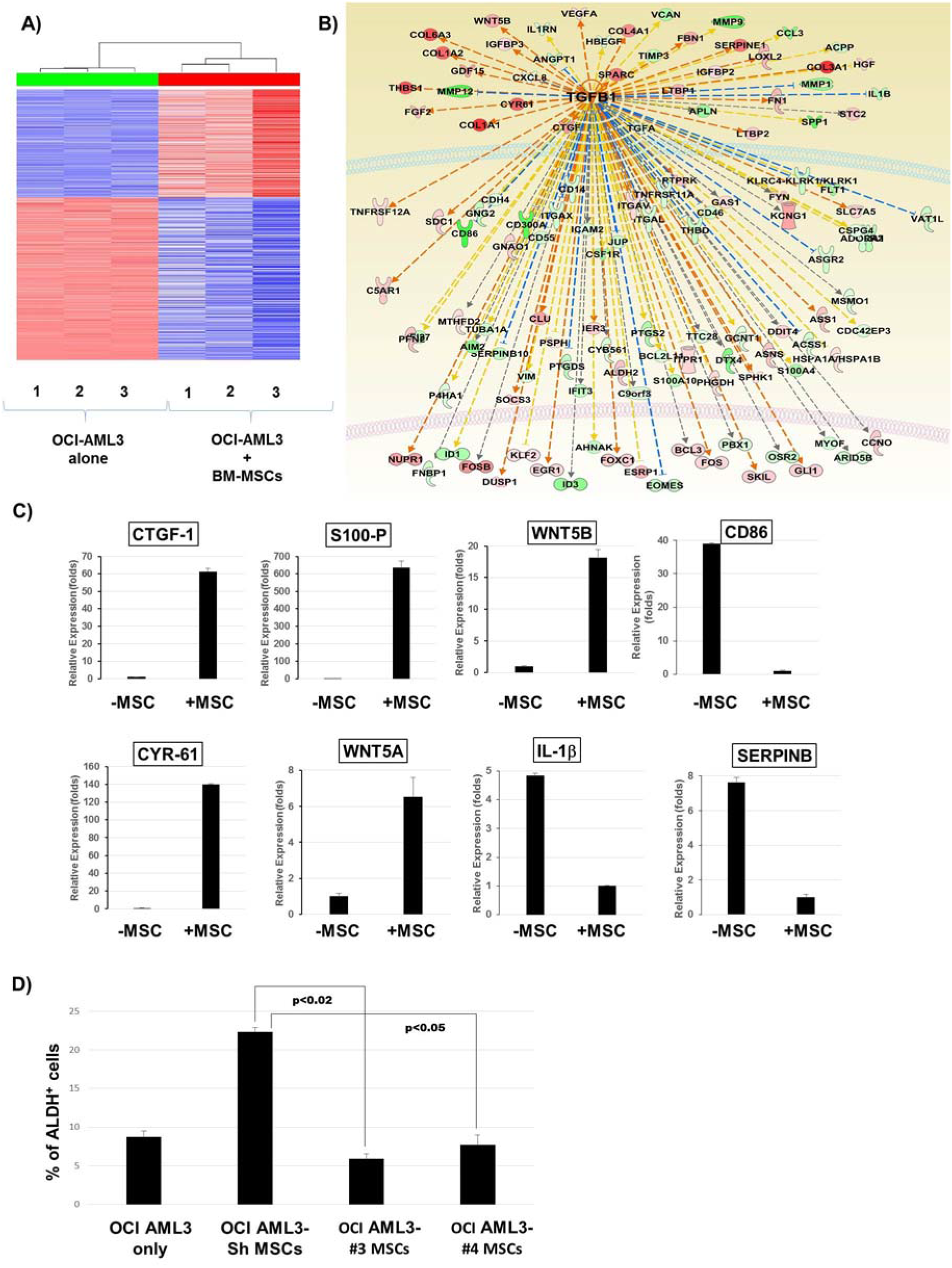
Activation of TGF-β1 gene signature in OCI-AML3 cells co-cultured with BM-MSCs. **(A, B)** OCI-AML3 cells were cultured with or without BM-MSC cells for 3 days. OCI-AML3 cells were FACS sorted to separate them from BM-MSCs and the gene expression analysis was performed by RNA sequencing. Samples were sequenced on the HiSeq Sequencing System. Sequence reads were mapped to human genomics (build hg19) with bowtie2 aligner using RSEM software. R software was used to compare the differential expression between MSC co-culture samples and OCI-AML3 controls. Genes with adjusted p values less than 0.05 and absolute fold changes larger than 2 were considered significant. Analysis of differentially expressed genes by Ingenuity^®^ pathway analysis tool revealed activation of TGF-β1-associated gene signature. **(C)** Real-time PCR analysis was performed to analyze the expression of indicated genes that are differentially regulated by TGF-β1 in AML cells co-cultured with BM-MSCs compared to OCI-AML3 controls. **(D)** TGF-β1 knockdown MSCs were generated by transfection with plasmid-containing lentiviruses and co-cultured with OCI-AML3 cells for 3 days. ALDH activity was measured by flow cytometry in OCI-AML3 cells cultured with TGF-β1 knockdown MSCs compared to OCI-AML3 cells cultured alone or with control BM-MSCs. Data are plotted as mean values with error bars representing standard error. For **C**, Mann-Whitney U-test or Student T-test was used. For **D**, one-way ANOVA with Tukey’s HSD post hoc test was used.

### Recombinant TGF-β1 induces ALDH activity in AML cells

It has been well-established that TGF-β1 signaling is involved in AML-BME interactions (5, 22). TGF-β is highly expressed in BM-MSCs and its expression is further enhanced in co-culture with leukemia cells (22). TGF-β has been shown to induce quiescence and stem-like phenotype in solid tumors as well as in leukemia (8). To test whether TGF-β1 induces ALDH activity in AML cells leading to a stem-like phenotype, we treated OCI-AML3 and HL60 cells with or without recombinant TGF-β1 (5 ng/mL) for 3 days and measured ALDH activity. Interestingly, treatment with recombinant TGF-β1 increased the percentage of ALDH^+^ cells from 30% ± 5% to 90% ± 3% and from 10% ± 4% to 25% ± 3% in OCI-AML3 and HL60 cells, respectively, suggesting that TGF-β1 directly regulates ALDH activity in AML cells (Fig. 4A, 4B).

**Figure 4.**
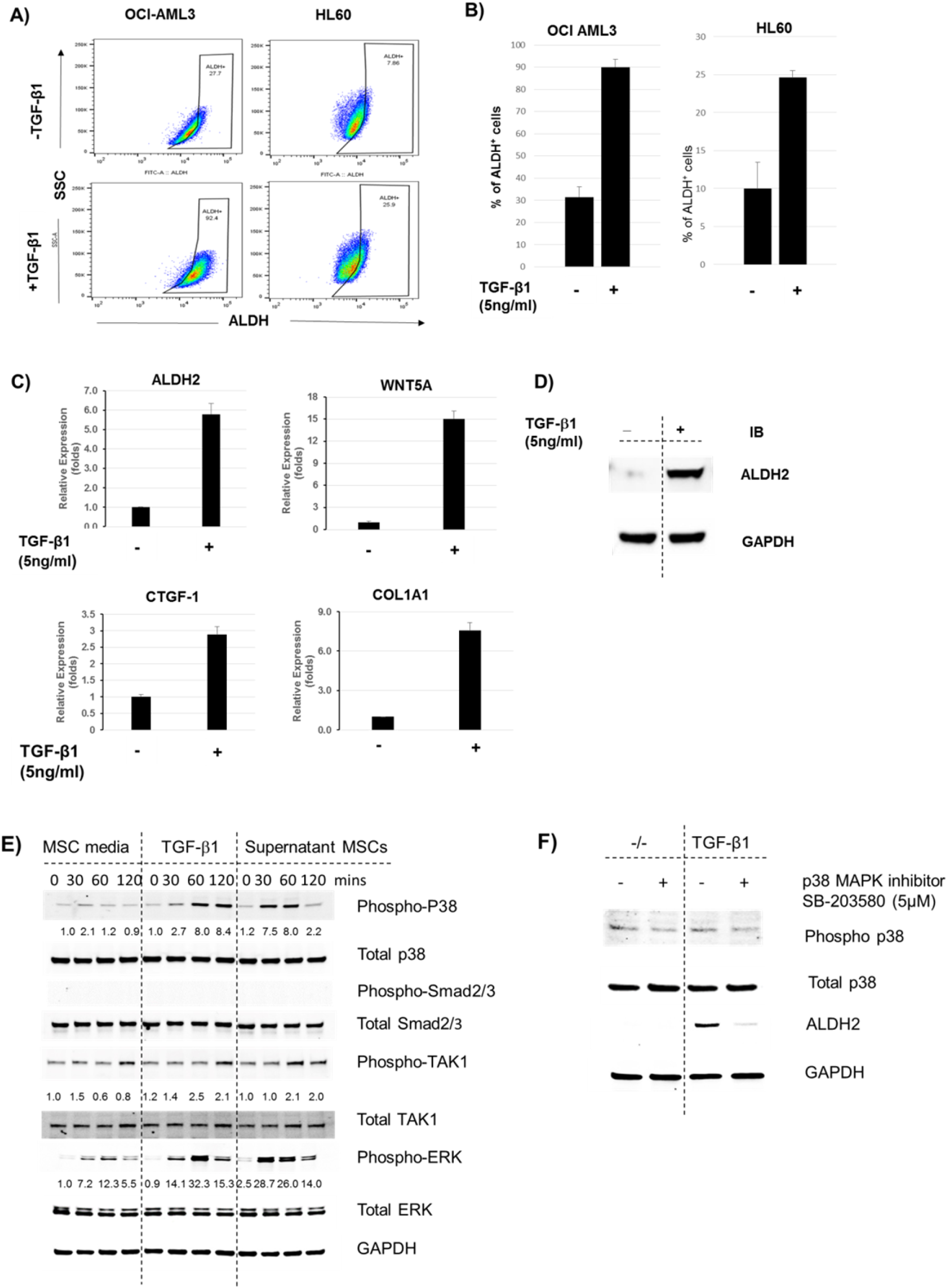
TGF-β1 non-canonical pathway induces ALDH2 and other downstream target expression in AML cells. **(A)** OCI-AML3 and HL60 cells were treated with recombinant TGF-β1 (5 ng/mL) for 3 days. Cells were stained with ALDEFLUOR^®^ and ALDH activity was measured. **(B)** Histogram representation of the percentage of ALDH^+^ cells in OCI-AML3 and HL60 AML cells treated with recombinant TGF-β1 in **A. (C)** OCI-AML3 cells were treated with recombinant TGF-β1 (5 ng/mL). RT-PCR was performed to analyze mRNA expression of downstream TGF-β targets, including WNT5A, CTGF1, and COL1A1. Relative fold increase values in gene expression were normalized to GAPDH. Data are plotted as mean values with error bars representing standard error (**D)** OCI-AML3 cells were treated with recombinant TGF-β1 (5 ng/mL) for 3 days. Protein lysates were harvested and western blotting was performed to analyze ALDH2 expression in treated cells compared to untreated controls. **(E)** OCI-AML3 cells were cultured in RPMI overnight. MSC cells were centrifuged and the MSC supernatant and TGF-β1 (5 ng/mL) were added to stimulate OCI-AML3 cells. Western blot analysis was performed at the indicated time points to analyze protein expression of downstream targets of the TGF-β canonical and non-canonical signaling pathways. **(F)** OCI-AML3 cells were treated with p38 MAPK inhibitor SB-203580 (5 μM) in the presence or absence of recombinant TGF-β1 (5 ng/mL). Protein lysates were harvested and p38 and ALDH2 expression was analyzed by western blotting. For **B** and **C**, Man-Whitney U-test and Student T-test were used, respectively (N=3).

To validate that TGF-β1 regulates downstream transcriptional activity in AML cells, we measured mRNA expression of TGF-β1 target genes in OCI-AML3 cells treated with recombinant TGF-β1. We found that TGF-β1 target genes were upregulated in cells treated with recombinant TGF-β1 compared to untreated controls (Fig. 4C). Interestingly, we also found that ALDH2, which was upregulated in AML cells upon co-culture with BM-MSCs, was also upregulated at the mRNA and protein levels in cells treated with recombinant TGF-β1 (Fig 4D, 4E).

### TGF-β non-canonical pathway is involved in stroma-induced ALDH activity in AML cells

TGF-β has been reported to alter downstream target gene expression through either its canonical pathway or its non-canonical pathway involving p38 and ERK (6, 23). To delineate the specific signaling pathway involved in TGF-β1-induced ALDH activity in AML cells, we treated OCI-AML3 cells with or without cell culture supernatants from BM-MSCs for 0, 30, 60, and 120 minutes and measured phosphorylation of transcription factors that mediate TGF-β1 canonical and non-canonical pathways by western blotting. Interestingly, we couldn’t detect any phosphorylation of Smad2 or Smad3 transcription factors in OCI-AML3 cells treated with supernatants derived from BM-MSCs. To confirm that BM-MSCs do not induce the canonical TGF-β pathway in AML cells, we measured phosphorylation of Smad2 and Smad3 in OCI-AML3 cells treated with recombinant TGF-β1 and still could not find any activity for these 2 proteins (Fig. 4E). Next, we tested phosphorylation of p38, which is activated by TGF-β through its non-canonical pathway. Interestingly, we found strong activity for phospho-p38 in OCI-AML3 cells treated with supernatants from BM-MSCs compared to cells treated with medium alone. We also found increased phosphorylation of ERK in OCI-AML3 cells treated with BM-MSC supernatants, suggesting activation of the Raf-MEK-ERK pathway in these cells. We validated this by treating OCI-AML3 cells with recombinant TGF-β1 and found a time-dependent increase in p38 phosphorylation (Fig. 4E).

p38 MAPK is a downstream target of the TGF-β non-canonical/non-Smad signaling pathway, which phosphorylates p38; p38 then activates other downstream signals leading to regulation of gene expression (22, 24, 25). To investigate the role of phospho-p38 in TGF-β1-mediated, stroma-induced ALDH2 expression in AML cells, OCI-AML3 cells were treated with or without p38 MAPK inhibitor SB-203580 (5 μM) and ALDH2 expression was measured by western blotting. We found that inhibition of p38 dramatically inhibited ALDH2 expression in OCI-AML3 cells, even in the presence of BM-MSC supernatant or recombinant TGF-β1 (Fig. 4F). This indicates that stromal cells induce ALDH2 expression in AML cells through non-canonical TGF-β/p38 MAPK pathway.

### p38 MAPK inhibition significantly inhibits ALDH activity in the presence of TGF-β or MSCs

To investigate the effect of p38 inhibition on ALDH2 activity, we treated OCI-AML3 cells with p38 MAPK inhibitor (5 μM) in the presence or absence of recombinant TGF-β1 (5 ng/mL) and measured ALDH activity. We found that p38 MAPK inhibition decreased the percentage of ALDH^+^ OCI-AML3 cells from 30% ± 5% to 11% ± 1%. Moreover, in the presence of recombinant TGF-β1, p38 MAPK inhibitor decreased the percentage of ALDH^+^ OCI-AML3 cells from 96% ± 1% to 22% ± 1% (Fig. 5A). Similarly, the addition of p38 MAPK inhibitor decreased the percentage of ALDH^+^ OCI-AML3 cells co-cultured with MSCs (Fig. 5B). These results solidify the notion that the non-canonical TGF-β pathway regulates ALDH activity in AML cells in the BME and that inhibition of p38 MAPK decreases TGF-β- or stroma-induced ALDH2 activity in AML cells.

**Figure 5.**
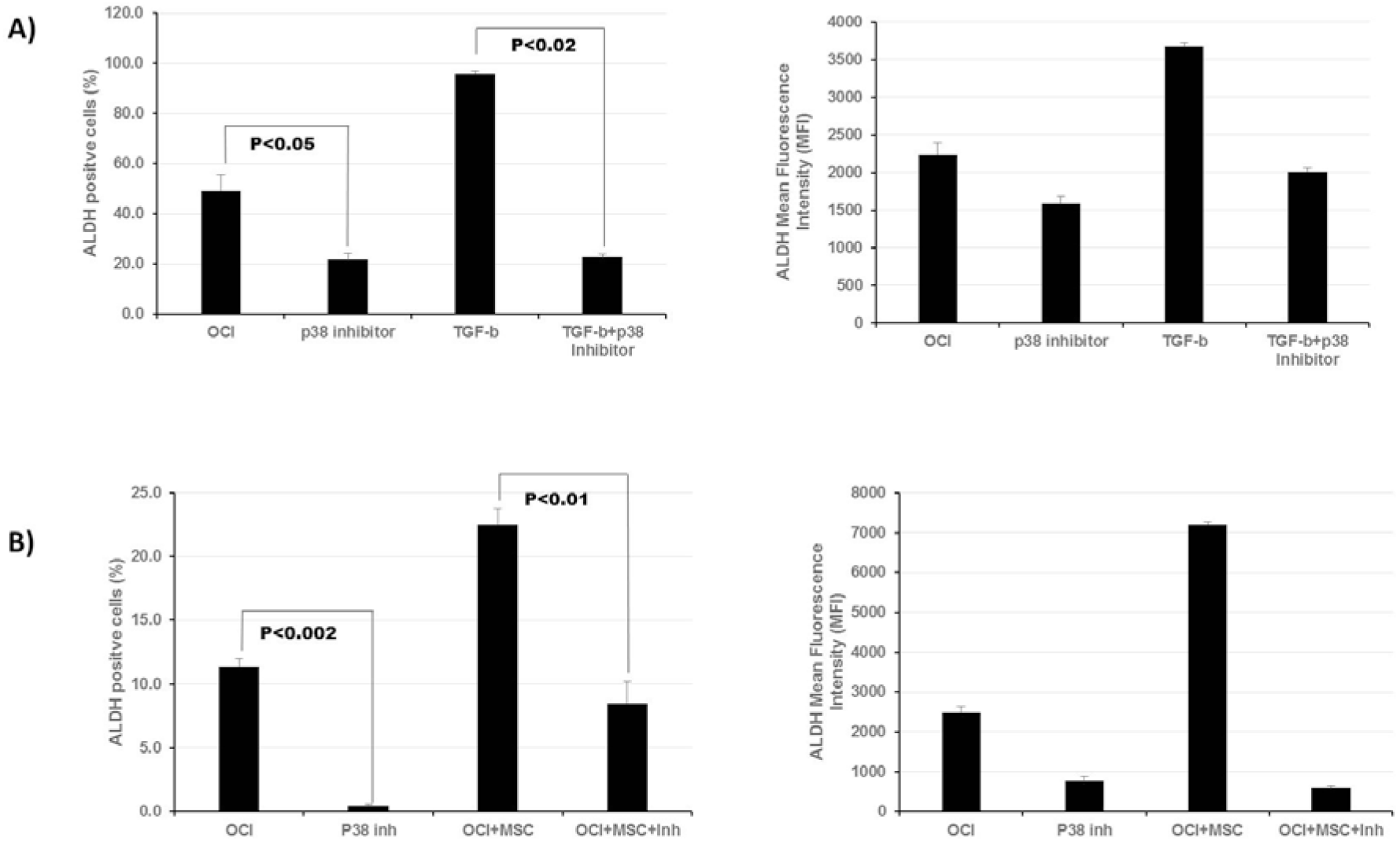
p38 MAPK inhibitor decreases ALDH2 expression in AML cells in the presence of TGF-β1 or stromal cells. **(A)**. OCI-AML3 cells were treated with p38 MAPK inhibitor SB-23580 (5 μM) in the presence or absence of recombinant TGF-β1 (5 ng/mL). ALDH activity was measured by flow cytometry using ALDEFLUOR^®^ assay in comparison to untreated controls. **(B**) OCI-AML3 were cultured with or without BM-MSCs and treated with p38 MAPK as in **A.** ALDH activity was measured in treated cells in comparison with untreated controls. Data are plotted as mean values with error bars representing standard error via one-way ANOVA with Tukey’s HSD post hoc test (N=3).

### ALDH2 inhibitors significantly decrease MSC-induced ALDH activity in AML cells

ALDH2 overexpression has been associated with several malignancies and diseases, leading to the development of specific inhibitors with potential therapeutic benefits (18, 26, 27). To validate that stroma-induced ALDH activity is mostly due to the ALDH2 isoform, we cultured OCI-AML3 cells and treated them with ALDH2 inhibitors diadzin or CVT-10216. Strikingly, treatment with ALDH2-specific inhibitors significantly inhibited ALDH activity in OCI-AML3 dose-dependently. The percentage of ALDH^+^ cells decreased from 29% ± 1% in untreated cells to 4% ± 0.5% in cells treated with 50 μM of diadzin. Similarly, the percentage of ALDH^+^ cells dropped from 16% ± 5% in untreated cells to 4% ± 0.2% in cells treated with 2 μM of CVT-10216 (Fig. 6A, 6B). Next, OCI-AML3 cells were treated with diadzin (5 μM) or CVT-10216 (1 μM) in the presence or absence of recombinant TGF-β1 (5 ng/mL) for 3 days. As expected, when treated with recombinant TGF-β1, the percentage of ALDH^+^ cells in OCI-AML3 cells increased from 23% ± 3% to 63% ± 2%. However, ALDH^+^ cells decreased from 63% ± 2% to 45% ± 2% when treated with diadzin, suggesting that diadzin inhibits ALDH activity even in the presence of TGF-β1. Similarly, we found an ~50% reduction in ALDH activity in OCI-AML3 cells treated with CVT-10216, even in the presence of recombinant TGF-β1 or BM-MSCs (Fig. 5B). This indicates that stroma-mediated TGF-β1-induced ALDH activity in AML cells can be decreased by ALDH2 inhibitors.

**Figure 6.**
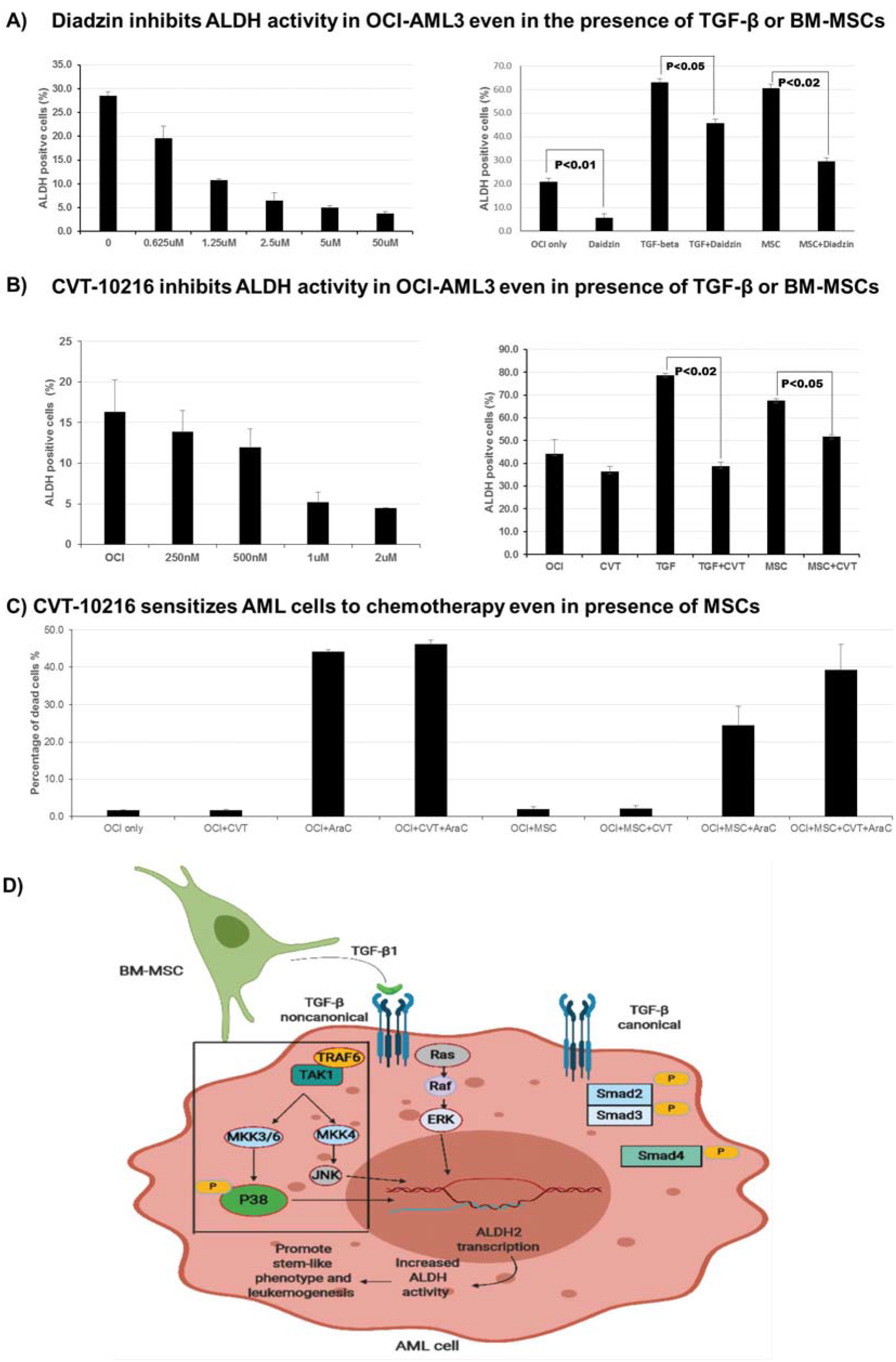
ALDH2 inhibitors decrease stroma-induced ALDH activity in AML cells and sensitize them to chemotherapy. **(A)** OCI-AML3 cells were cultured and treated with increasing concentrations of ALDH2 inhibitor diadzin. ALDH activity was measured by ALDEFLUOR^®^ assay using flow cytometry. OCI-AML3 cells were co-cultured with MSCs (200 000 cells/well density) and treated with diadzin (5 μM) for 3 days in the presence or absence of recombinant TGF-β1 (5 ng/mL). ALDH activity was measured as described above. ALDH activity in OCI-AML3 cells co-cultured with MSCs (200 000 cells/well density). **(B)** The same experiments were performed as in **A**, using CVT-10216 (1 μM) instead of diadzin. **(C)** OCI-AML3 cells were cultured alone or with MSCs (100 000 cells/well density) and treated with cytarabine (2.5 μM) alone or in combination with CVT-10216 (2 μM) for 48 hours. Cell samples were stained with Annexin V and analyzed by flow cytometry to measure treatment-induced cell death. Data are plotted as mean values with error bars representing standard error. **(D)** Simplified schematic representation of the interaction between stromal cells and AML cells leading to ALDH2 expression. BM-MSCs secrete TGF-β1, which induces ALDH2 expression in AML cells through the non-canonical/p38-dependent pathway, thereby promoting leukemogenesis. For **A**, **B**, and **C** one-way ANOVA with Tukey’s HSD post hoc test (N=3).

### ALDH2 inhibitors sensitize AML cells to standard chemotherapy

To evaluate the therapeutic significance of ALDH2 inhibition in AML, we tested the effect of CVT-10216 in combination with standard chemotherapy on AML cell death. We cultured OCI-AML3 cells with or without BM-MSCs and treated them with cytarabine (2.5 μM) alone or in combination with CVT-10216 (2 μM) for 48 hours. The cells were stained with Annexin V and DAPI and analyzed by flow cytometry to measure treatment-induced apoptosis and cell death. Interestingly, treatment with CVT-10216 significantly improved cytarabine-induced cell death in AML cells. The combination of CVT-10216 and cytarabine induced cell death in 39% ± 6% of OCI-AML3 cells compared to 24% ± 5% cells treated with cytarabine alone (Fig. 6C.). Our results suggest ALDH2 is a potential therapeutic target in AML and that CVT-10216 sensitizes AML cells to conventional chemotherapy.

## Discussion

In the present study, we demonstrate that BM-MSCs contribute to AML progression and chemo-resistance by inducing an ALDH^+^ stem-like phenotype in AML cells. Importantly, through our in-depth gene expression analysis of all 19 ALDH isoforms, we identified ALDH2 as the isoform which is differentially expressed and primarily responsible for the increased ALDH activity in the presence of stroma, rendering it a major contributor to AML stem-like phenotype and consequent chemotherapy resistance.

Recent studies have demonstrated that TGF-β is an important mediator in AML-BME interactions and is involved in conferring AML chemotherapy resistance (8, 28). Our results are not unlike findings reported in the literature, as we have shown upregulation of the TGF-β1 gene signature in AML cells in the presence of stromal cells. We also show that downstream targets of TGF-β signaling, including ALDH2, are differentially expressed in AML cells in the presence of MSCs. Moreover, exposure of AML cells to recombinant TGF-β1 induced ALDH activity through ALDH2 expression, but knockdown of TGF-β1 in BM-MSCs led to a decrease in ALDH activity, indicating that TGF-β1 secreted by BM-MSCs induces a stem-like phenotype in AML cells. Additionally, through protein expression analysis, we delineated the specific signaling pathway involved in the aforementioned TGF-β1-mediated AML-BME interaction. We showed that TGF-β1 exerts its effect through a non-canonical/p38-dependent signaling pathway. Hence, our work uncovers a clear link between TGF-β1 secreted by MSCs and the acquisition of a stem-like phenotype of AML cells through ALDH2 overexpression; it also establishes the specific signaling mechanism involved in this interaction.

ALDH plays a prominent role in several diseases and malignancies (18, 29), and specific ALDH inhibitors with potentially anti-tumor effects have been developed (17, 26, 30–32). In this study, we examined the role of ALDH2 inhibitors in counteracting the stem-like phenotype and chemotherapy resistance acquired by AML cells in the presence of stromal cells. Interestingly, we reveal that specific ALDH2 inhibitors significantly inhibit ALDH activity in a dose-dependent manner and sensitize AML cells to conventional treatment, even in the presence of MSCs. Although further studies are needed to confirm this, our preliminary results identify stroma-induced ALDH2 as a novel target for treatment of refractory AML in combination with standard chemotherapy.

Our study has some potential limitations. We show promising results regarding ALDH2 inhibition as a strategy to counteract AML chemotherapy resistance; however, we have not validated these findings in vivo. Additionally, ALDH2 inhibitors tested significantly reduced, but did not completely inhibit, ALDH activity in AML cells. It remains unclear whether other ALDH isoforms contribute to ALDH activity or the ALDH2 inhibitors we used are not sufficiently potent and efficacious. Moreover, there could be other potential mechanisms of stem-cell phenotype acquisition in AML beyond the scope of our study. Despite the above limitations, our work remains a valuable addition to the literature on AML-microenvironment interactions as it provides a coherent and comprehensive framework for a better understanding of the role of stromal cells in AML chemotherapy resistance and identifies several potential clinically relevant therapeutic targets to consider.

Our work sets the foundation for future studies to further investigate the therapeutic applications of targeting mediators of the AML-stroma interactions that we identified in this report. Further studies are needed to examine the effect of ALDH2 inhibitors in combination with standard chemotherapy in vivo, setting the framework for future phase 1/phase 2 clinical trials assessing the therapeutic efficacy of such inhibitors for refractory AML.

In summary, BM stromal cells induce ALDH activity in AML cells through increased expression of the ALDH2 isoform. BM-MSCs secrete TGF-β1, which exerts its effect through a non-canonical/p38-dependent signaling mechanism, leading to a stem-like phenotype in AML cells. Inhibition of downstream targets of this pathway, such as p38 MAPK, inhibits ALDH activity in AML cells. ALDH2 inhibition sensitizes AML cells to standard chemotherapy in vitro, providing a potential novel therapeutic strategy to overcome AML chemotherapy resistance.

## Author Contributions

BY and FE conducted experiments, analyzed the data and wrote the manuscript. SL and VR performed experiments. YY performed all statistical tests and analyses. ES and MK analyzed and interpreted the data. MA planned the study, interpreted the results, and edited the manuscript. VLB conceptualized the study, planned all experiments, analyzed and interpreted the results, and edited the manuscript. All authors read the final manuscript and approved it.

## List of Abbreviations

1. ALDH: Aldehyde dehydrogenase
2. AML: Acute myeloid leukemia
3. BM: Bone marrow
4. BM-MSC: Bone marrow derived mesenchymal stromal cells
5. FBS: Fetal bovine serum
6. MSC: Mesenchymal stromal cells
7. PBS: Phosphate buffered saline
8. PBS-T: 0.05% Tween 20 in phosphate buffered saline
9. PVDF: Polyvinylidene fluoride
10. TGF-β: Transforming growth factor-β

## References

1. Role of the bone marrow microenvironment in CML and AML. Bonekey Rep. 2014 Jan 15;3:504.

2. Chang YT, Hernandez D, Alonso S, Gao M, Su M, Ghiaur G, et al. Role of CYP3A4 in bone marrow microenvironment-mediated protection of FLT3/ITD AML from tyrosine kinase inhibitors. Blood Adv. 2019 Mar 26;3(6):908–16.

3. Zeng Z, Shi YX, Tsao T, Qiu Y, Kornblau SM, Baggerly KA, et al. Targeting of mTORC1/2 by the mTOR kinase inhibitor PP242 induces apoptosis in AML cells under conditions mimicking the bone marrow microenvironment. Blood. 2012 Sep 27;120(13):2679–89.

4. Viswanathan VS, Ryan MJ, Dhruv HD, Gill S, Eichhoff OM, Seashore-Ludlow B, et al. Dependency of a therapy-resistant state of cancer cells on a lipid peroxidase pathway. Nature. 2017 Jul 27;547(7664):453–7.

5. Tabe Y, Shi YX, Zeng Z, Jin L, Shikami M, Hatanaka Y, et al. TGF-beta-Neutralizing Antibody 1D11 Enhances Cytarabine-Induced Apoptosis in AML Cells in the Bone Marrow Microenvironment. PLoS One. 2013;8(6):e62785.

6. Baarsma HA, Menzen MH, Halayko AJ, Meurs H, Kerstjens HA, Gosens R. beta-Catenin signaling is required for TGF-beta1-induced extracellular matrix production by airway smooth muscle cells. Am J Physiol Lung Cell Mol Physiol. 2011 Dec;301(6):L956–65.

7. Baarsma HA, Spanjer AI, Haitsma G, Engelbertink LH, Meurs H, Jonker MR, et al. Activation of WNT/beta-catenin signaling in pulmonary fibroblasts by TGF-beta(1) is increased in chronic obstructive pulmonary disease. PLoS One. 2011;6(9):e25450.

8. Caja L, Bertran E, Campbell J, Fausto N, Fabregat I. The transforming growth factor-beta (TGF-beta) mediates acquisition of a mesenchymal stem cell-like phenotype in human liver cells. J Cell Physiol. 2011 May;226(5):1214–23.

9. Yadav P, Shankar BS. Radio resistance in breast cancer cells is mediated through TGF-beta signalling, hybrid epithelial-mesenchymal phenotype and cancer stem cells. Biomed Pharmacother. 2019 Mar;111:119–30.

10. Borges L, Oliveira VKP, Baik J, Bendall SC, Perlingeiro RCR. Serial transplantation reveals a critical role for endoglin in hematopoietic stem cell quiescence. Blood. 2019 Feb 14;133(7):688–96.

11. Brandes AA, Carpentier AF, Kesari S, Sepulveda-Sanchez JM, Wheeler HR, Chinot O, et al. A Phase II randomized study of galunisertib monotherapy or galunisertib plus lomustine compared with lomustine monotherapy in patients with recurrent glioblastoma. Neuro Oncol. 2016 Aug;18(8):1146–56.

12. Garaycoechea JI, Crossan GP, Langevin F, Daly M, Arends MJ, Patel KJ. Genotoxic consequences of endogenous aldehydes on mouse haematopoietic stem cell function. Nature. 2012 Sep 27;489(7417):571–5.

13. Storms RW, Trujillo AP, Springer JB, Shah L, Colvin OM, Ludeman SM, et al. Isolation of primitive human hematopoietic progenitors on the basis of aldehyde dehydrogenase activity. Proc Natl Acad Sci U S A. 1999 Aug 3;96(16):9118–23.

14. Cheung AM, Wan TS, Leung JC, Chan LY, Huang H, Kwong YL, et al. Aldehyde dehydrogenase activity in leukemic blasts defines a subgroup of acute myeloid leukemia with adverse prognosis and superior NOD/SCID engrafting potential. Leukemia. 2007 Jul;21(7):1423–30.

15. Ran D, Schubert M, Pietsch L, Taubert I, Wuchter P, Eckstein V, et al. Aldehyde dehydrogenase activity among primary leukemia cells is associated with stem cell features and correlates with adverse clinical outcomes. Exp Hematol. 2009 Dec;37(12):1423–34.

16. Yang L, Chen WM, Dao FT, Zhang YH, Wang YZ, Chang Y, et al. High aldehyde dehydrogenase activity at diagnosis predicts relapse in patients with t(8;21) acute myeloid leukemia. Cancer Med. 2019 Sep;8(12):5459–67.

17. Chen CH, Ferreira JC, Gross ER, Mochly-Rosen D. Targeting aldehyde dehydrogenase 2: new therapeutic opportunities. Physiol Rev. 2014 Jan;94(1):1–34.

18. Koppaka V, Thompson DC, Chen Y, Ellermann M, Nicolaou KC, Juvonen RO, et al. Aldehyde dehydrogenase inhibitors: a comprehensive review of the pharmacology, mechanism of action, substrate specificity, and clinical application. Pharmacol Rev. 2012 Jul;64(3):520–39.

19. Battula VL, Le PM, Sun JC, Nguyen K, Yuan B, Zhou X, et al. AML-induced osteogenic differentiation in mesenchymal stromal cells supports leukemia growth. JCI Insight. 2017 Jul 6;2(13).

20. Battula VL, Chen Y, Cabreira Mda G, Ruvolo V, Wang Z, Ma W, et al. Connective tissue growth factor regulates adipocyte differentiation of mesenchymal stromal cells and facilitates leukemia bone marrow engraftment. Blood. 2013 Jul 18;122(3):357–66.

21. Yang X, Sexauer A, Levis M. Bone marrow stroma-mediated resistance to FLT3 inhibitors in FLT3-ITD AML is mediated by persistent activation of extracellular regulated kinase. Br J Haematol. 2014 Jan;164(1):61–72.

22. Koschmieder S, Hofmann WK, Kunert J, Wagner S, Ballas K, Seipelt G, et al. TGF beta-induced SMAD2 phosphorylation predicts inhibition of thymidine incorporation in CD34+ cells from healthy donors, but not from patients with AML after MDS. Leukemia. 2001 Jun;15(6):942–9.

23. Eger A, Stockinger A, Park J, Langkopf E, Mikula M, Gotzmann J, et al. beta-Catenin and TGFbeta signalling cooperate to maintain a mesenchymal phenotype after FosER-induced epithelial to mesenchymal transition. Oncogene. 2004 Apr 8;23(15):2672–80.

24. Abrigo J, Campos F, Simon F, Riedel C, Cabrera D, Vilos C, et al. TGF-beta requires the activation of canonical and non-canonical signalling pathways to induce skeletal muscle atrophy. Biol Chem. 2018 Feb 23;399(3):253–64.

25. Zhang Y, Zeng Y, Liu T, Du W, Zhu J, Liu Z, et al. The canonical TGF-beta/Smad signalling pathway is involved in PD-L1-induced primary resistance to EGFR-TKIs in EGFR-mutant non-small-cell lung cancer. Respir Res. 2019 Jul 22;20(1):164.

26. Wang F, Zhai S, Liu X, Li L, Wu S, Dou QP, et al. A novel dithiocarbamate analogue with potentially decreased ALDH inhibition has copper-dependent proteasome-inhibitory and apoptosis-inducing activity in human breast cancer cells. Cancer Lett. 2011 Jan 1;300(1):87–95.

27. Zhang Y, Ren J. ALDH2 in alcoholic heart diseases: molecular mechanism and clinical implications. Pharmacol Ther. 2011 Oct;132(1):86–95.

28. Dourado KMC, Baik J, Oliveira VKP, Beltrame M, Yamamoto A, Theuer CP, et al. Endoglin: a novel target for therapeutic intervention in acute leukemias revealed in xenograft mouse models. Blood. 2017 May 4;129(18):2526–36.

29. Lowe ED, Gao GY, Johnson LN, Keung WM. Structure of daidzin, a naturally occurring anti-alcohol-addiction agent, in complex with human mitochondrial aldehyde dehydrogenase. J Med Chem. 2008 Aug 14;51(15):4482–7.

30. Buchman CD, Hurley TD. Inhibition of the Aldehyde Dehydrogenase 1/2 Family by Psoralen and Coumarin Derivatives. J Med Chem. 2017 Mar 23;60(6):2439–55.

31. Buchman CD, Mahalingan KK, Hurley TD. Discovery of a series of aromatic lactones as ALDH1/2-directed inhibitors. Chem Biol Interact. 2015 Jun 5;234:38–44.

32. Kiyuna LA, Albuquerque RPE, Chen CH, Mochly-Rosen D, Ferreira JCB. Targeting mitochondrial dysfunction and oxidative stress in heart failure: Challenges and opportunities. Free Radic Biol Med. 2018 Dec;129:155–68.

